# Vnd and En are expressed in orthogonal stripes and act in a brief competence window to combinatorially specify NB7-1 and its early lineage

**DOI:** 10.1101/2025.09.04.674256

**Authors:** Nathan Anderson, Sen-Lin Lai, Chris Q. Doe

## Abstract

Understanding how neuronal diversity is generated is a major goal of neuroscience. Here we characterize the first step in generating neuronal diversity in the *Drosophila* embryo: spatial transcription factors (STFs) expressed in orthogonal rows and columns of neural progenitors. These factors give spatial identity to neural progenitors (neuroblasts, NBs), and are highly conserved in mammals. Here we investigate the roles of Engrailed (En+; posterior row) and Vnd+ (medial column) in specifying the well-characterized progenitor: neuroblast 7-1 (NB7-1). We show that NB7-1 is located at the intersection of Vnd and En, and we identify NB7-1 using a newly characterized gene, *fd4*, that we show is specifically expressed in NB7-1 and its progeny, giving us a specific assay for NB7-1 identity. We show that En and Vnd are both required for Fd4 expression, and that Vnd and En co-expression is sufficient to induce ectopic Fd4 expression in other NBs and their lineages. Finally, we show that NBs gradually lose competence to respond to En or Vnd. We conclude that En and Vnd are STFs that act combinatorially to specify the identity of an individual progenitor, NB7-1.

## Introduction

Neuronal diversity is required for proper circuit formation and behavior. Understanding how neuronal diversity is generated can help clarify the mechanisms of circuit assembly, as well as help design more efficient neuronal reprogramming necessary for treating brain injury and disease. In both mammals and *Drosophila*, there are two major steps in generating neuronal diversity: spatial and temporal patterning (Sagner et al., 2021; Skeath and Thor, 2003). Here we focus on spatial patterning.

In mammals, spatial patterning creates molecularly distinct pools of neural stem cells (NSCs), best characterized in the spinal cord. Here, opposing gradients of Bmp (high dorsally) and Sonic hedge hog (Shh) generate 11 molecularly distinct progenitor pools, which are committed to generating a distinct family of neurons and glia (Briscoe et al., 2000; Sagner et al., 2021).

In *Drosophila*, spatial patterning also generates progenitor diversity, which has been best characterized in the ventral nerve cord (VNC), a structure similar to the spinal cord (Skeath and Thor, 2003). The neuroectoderm expresses proneural genes that promote delamination of the nascent progenitors (neuroblasts, NBs in *Drosophila*) (Skeath et al., 1994; Skeath and Carroll, 1992). The initial 10 NBs in each hemisegment are arranged in rows (1, 3, 5, 7) and columns (ventral, intermediate, dorsal)(see Figure 1F). Additional rows and columns are added later. Genes expressed in rows or columns have been termed “spatial transcription factors (STFs)” when they are required for specifying NBs within rows or columns (Chu et al., 1998; McDonald et al., 1998; Weiss et al., 1998), and additional row and column factors are termed candidate STFs. To date there are three columnar STFs: the homeodomain factors Ventral nervous system defective (Vnd), Intermediate nervous system defective (Ind) and Muscle segment homeobox (Msh; Flybase: Drop) (Chu et al., 1998; McDonald et al., 1998; Weiss et al., 1998). Indeed, previous work showed that each of the columnar TFs function to specify the formation and specification of “columnar identity” within the VNC; for example, misexpression of the ventral factor Vnd can transform intermediate columnar-specific molecular markers with ectopic ventral column markers (Chu et al., 1998; McDonald et al., 1998), and misexpression of Ind can transform the lateral column into an expanded intermediate column (Weiss et al., 1998). These columnar STFs are strongly conserved in mammals where each ortholog (Nkx2.2, Gsx, and Msx) is also expressed in adjacent columns in the spinal cord (Weiss et al., 1998).

**Figure 1.**
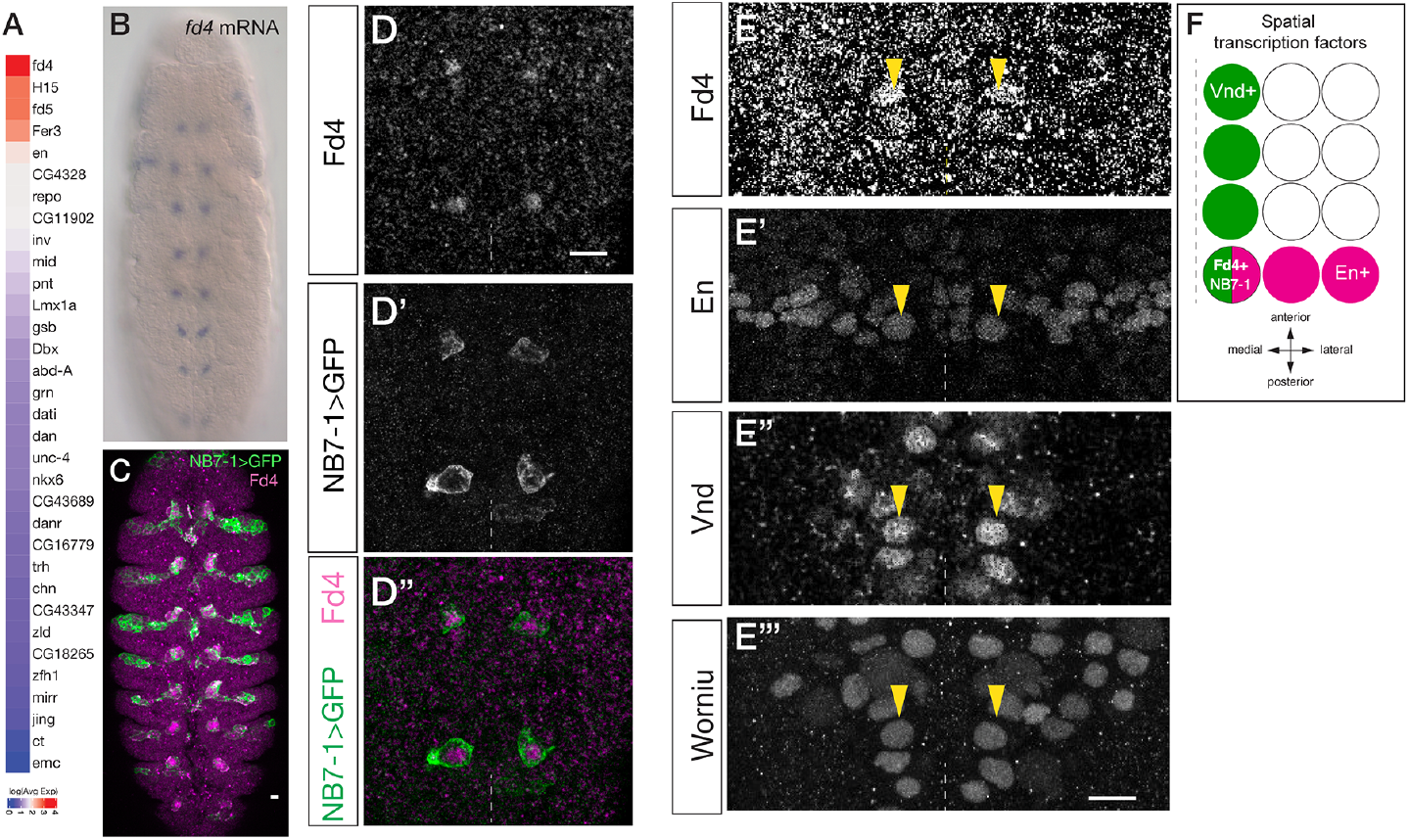
Fd4 is expressed in NB7-1, and its lineage, located at the junction of Vnd and En spatial factors. (A) Heatmap of log fold-change of the scaled average expression of transcription factors between NB7-1 lineage and other lineages. (B) *fd4* RNA is detected in a single NB lineage in each hemisegment. (C) *NB7-1-Gal4 UAS-GFP* (NB7-1>GFP), green; Fd4 protein, magenta. Stage 12 embryo, ventral view. Scale bar, 10μm. (D-D’’) Fd4 protein is co-expressed with NB7-1-Gal4. Stage 10 embryo, ventral view; two segments shown; midline, dashed line; scale bar, 10μm. (E-E’’’) Fd4 protein is co-expressed with Wor (NB marker), En and Vnd. Yellow arrowheads, Fd4+ NB. One segment shown; midline, dashed line; Stage 10. (F) Summary. Fd4 is expressed at the overlap between Vnd (column) and En (row).

Similarly, several row spatial factors are necessary and sufficient to specify “row identity” of new-born NBs: the signaling ligand Wingless expression in row 5 is required for formation and specification of NBs in the adjacent rows 4 and 6 (Chu-LaGraff and Doe, 1993), whereas Gsb is necessary and sufficient to specify row 5 identity (Bhat, 1996; Skeath et al., 1995). There are also candidate STFs: Mirror and Engrailed (En) from anterior to posterior (Chu-LaGraff and Doe, 1993; O’Farrell et al., 1985; Skeath et al., 1995).

These studies established the concept of spatial patterning via orthogonally expressed homeodomain STFs, but many open questions remain. Are there more STFs than have been characterized to date? Do row and column STFs act combinatorially to specify one or a few progenitors? When, if ever, do progenitors lose competence to respond to spatial patterning genes? Here we address all of these questions, using new NB and lineage markers.

## Methods

### Fly stocks

Flies were maintained in an incubator at 25C. The chromosomes and insertion sites of transgenes (if known) are shown next to genotypes. The following fly stocks were used: *en-gal4* (Schmid et al., 1999), *grainy head (grn)-gal4* (RRID:BDSC_76172), *NB7-1-gal4*^*KZ*^ (II) called *NB7-1-gal4* (Seroka and Doe, 2019), *scabrous (sca)-gal4* (Cheesman et al., 2004), *vnd-T2A-gal4* (called *vnd-gal4*; this work), *worniu (wor)-gal4* (Albertson and Doe, 2003), *10xUAS-IVS-myr::sfGFP* (*UAS-Gfp*) (RRID:BDSC_62127), *UAS-lacZ* (II) (RRID:BDSC_1776), *UAS-en* (Tabata et al., 1995), *UAS-vnd* (Chu et al., 1998), *vnd[A]* mutant (flybase.org/reports/FBal0298950.html) and *Df(2R)en[E]* which removes both redundant *en* and *inv* loci (RRID:BDSC_2216).

### Immunostaining and imaging

We performed overnight embryo collections onto yeasted apple juice agar caps. Standard methods were used to fix and stain embryos (Grosskortenhaus et al., 2005). Primary antibodies used were: rat anti-Dpn (1:20, Doe lab), mouse anti-En, 5mg/mL (Developmental Studies Hybridoma Bank (DHSB; Iowa City, IA), RRID:AB_528224), mouse anti-Eve[2B8], 5mg/mL, (DSHB, RRID_AB:528230), guinea pig anti-Fd4, 5mg/mL (Doe lab, this work), rabbit anti-Hb (1:300) (Tran and Doe, 2008), guinea pig anti-Runt (1:1000) (Sullivan et al., 2019), mouse anti-Wg, 5mg/mL (DSHB, RRID_AB:528512), rabbit anti-Wor, 1:1000 (Doe lab), rat anti-Zfh2 (1:300) (Tran et al., 2010). Secondary antibodies used were: DyLight 405, AlexaFluor 488, AlexaFluor 555, Rhodamine Red^TM^-X (RRX), or Alexa Fluor 647-conjugated AffiniPure^TM^ donkey anti-IgG (Jackson ImmunoResearch, West Grove, PA, USA). The samples were mounted in 90% glycerol with Vectashield (Vector Laboratories, Burlingame, CA). Images were captured with a Zeiss LSM 900 confocal microscope with a *z*-resolution of 0.5 μm. Images were processed using the open-source software FIJI, or using Imaris (Oxford Instruments plc, UK). Figures were assembled in Adobe Illustrator (Adobe, San Jose, CA).

### Identification of U1-U5 neurons

The five Even-skipped (Eve)+ U neurons were be individually identified using the following molecular markers and cell body position. U1, Hb+; U2, Hb+ Zfh2+; U3, Zfh2+; U4, Runt+ anterior-medial; U5, Runt+ posterior-lateral (Cleary and Doe, 2006; Isshiki et al., 2001; Pearson and Doe, 2003).

### Newly generated genetic tools

To make vnd-T2A-gal4 flies, T2A-Gal4 was integrated into the C-terminus of *vnd* open reading frame with CRISPR using the pHD-DsRed (Addgene plasmid #51434; http://n2t.net/addgene:51434; RRID:Addgene_51434) plasmid. This plasmid also included 1Kb of homologous arms both upstream and downstream of the last codon, with the stop codon removed, for homology directed repair (HDR). Two gRNAs CCCACGCCCAGAGGGCCGCC and TGCCAGTTCCTAGCAATATT were used and cloned into pCFD5 (Addgene plasmid #73914; http://n2t.net/addgene:49411; RRID:Addgene_73914). The gRNA and HDR constructs were co-injected into the *yw*;*nos-cas9* (II-attP40) flies by BestGene (Chino Hills, CA, USA). Six independent transformants were obtained and their expression patterns were verified.

The Fd4 antibody was generated with full length protein immunized in guinea pigs by GenScript (Piscataway, NJ, USA) and its detailed expression pattern and function will be reported separately (Lai and Doe, in preparation); here we use it simply as a marker for NB7-1.

### Statistics

All analyses were conducted in Prism. *P*-values are reported in the figure legends. Plots display n.s. = not significant, * is *P* < 0.05, and ** is *P* < 0.01.

## Results

### Fd4 is specifically expressed in NB7-1 and its lineage

To explore the mechanisms by which NBs establish and retain their unique identity, we focused on the NB7-1. We previously generated a single cell RNA sequencing library which contains NB7-1 lineage and other NB lineages (Seroka et al., 2022). We identified the gene *fd4* (FlyBase: *fd96Ca*) as the most enriched transcription factor in NB7-1 lineage (Figure 1A), suggesting Fd4 could have a NB7-1-specific expression and function. *fd4* RNA was expressed in a segmentally repeated cluster of cells adjacent to the midline, corresponding to the region where the NB7-1 lineage is located (Fig. 1B; https://tinyurl.com/fd4infly). We generated an antibody recognizing Fd4 protein (see methods), and found Fd4 is specifically detected in cells expressing NB7-1-gal4 UAS-GFP+, thereby validating its specific expression in the NB7-1 lineage (Fig. 1C,D).

To determine the STFs that NB7-1 expresses, we stained embryos for Vnd (medial column), En (posterior row), and Fd4 (NB7-1). We found that the Fd4+ NB7-1 was the sole NB that expressed both Vnd and En (Figure 1E). We confirmed that only the En/Vnd double-positive NB expressed Fd4 (Figure 1E, and summarized in 1F). For a more detailed analysis of Fd4 expression and function, see SLL and CQD (in preparation); here we used Fd4 solely as a specific marker for NB7-1. We conclude that Fd4 is specifically expressed in NB7-1, which forms at the junction of Vnd and En expression domains.

### The spatial transcription factors Vnd and En are both required for Fd4 expression

Fd4 was expressed in the only NB that is both En+ and Vnd+, raising the question of whether Vnd and En function together to drive expression of Fd4 in NB7-1 identity. In control embryos, NB7-1 expressed Fd4, the NB marker Worniu (Wor), and the spatial factors Vnd-GFP and En (Figure 2A). Note that a Vnd antibody does not exist, so we used a GFP-tagged Vnd to report on Vnd expression. In contrast, removal of the redundant, tandem *en*/*invected* genes resulted in complete loss of Fd4 expression without altering Wor or Vnd-GFP expression (Figure 2B-B’’’). Similarly, removal of Vnd in a homozygous null background resulted in complete loss of Fd4 expression, without altering Wor or En expression (Figure 2C-C’’’). We conclude that the Vnd and En STFs are both required to drive expression of Fd4 in NB7-1 (Figure 2D).

**Figure 2.**
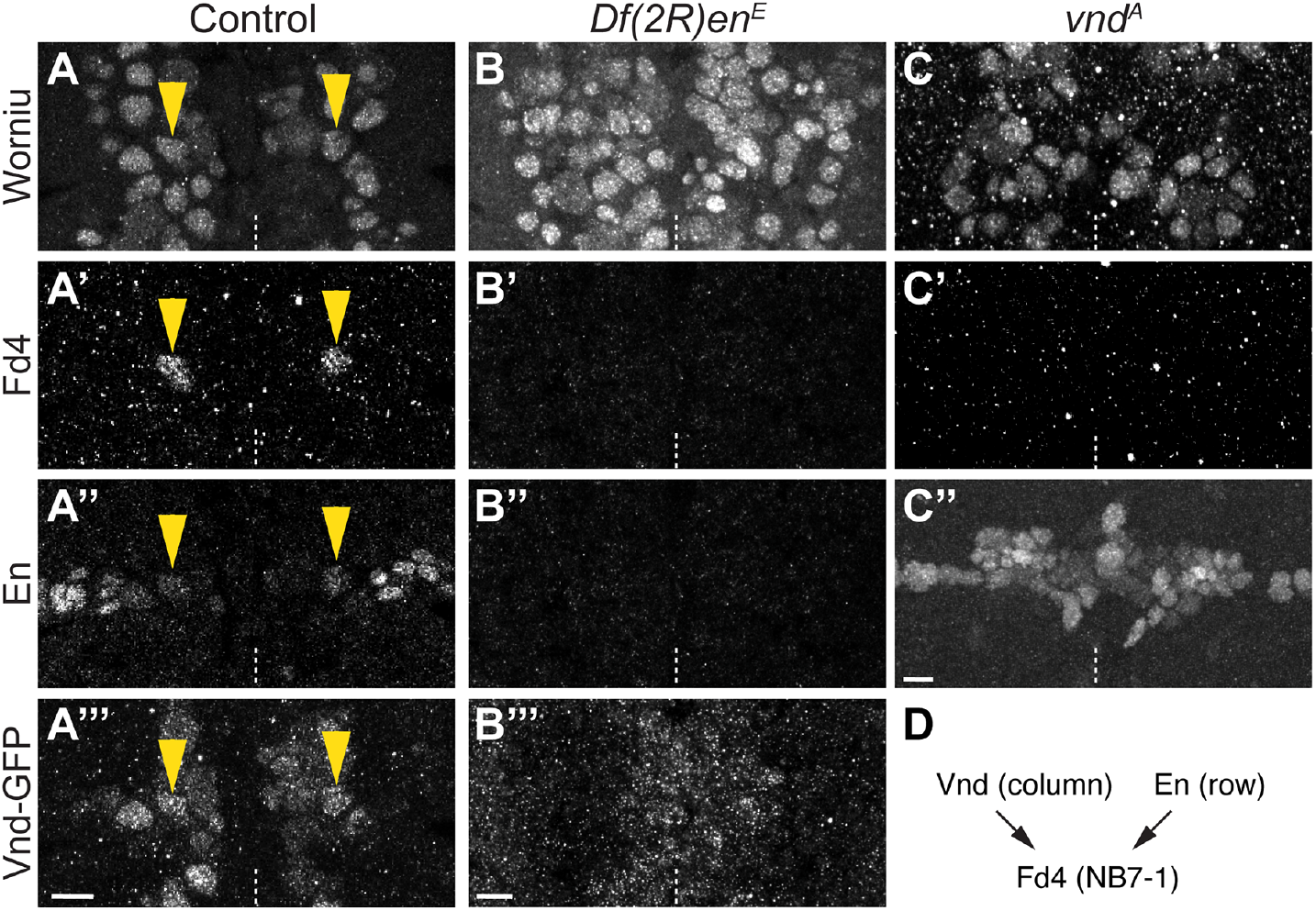
Both Vnd and En are required for Fd4 expression. (A) Control. Worniu (NB marker), Fd4, En, and Vnd. For this and other panels; in stage 11 embryo; one segment shown, ventral view; midline, dashed line; yellow arrowhead, Fd4+ NB. Scale bar 10μm. (B) Loss of En results in complete loss of Fd4, without affecting Worniu or Vnd-GFP. (C) Loss of Vnd results in complete loss of Fd4, without affecting Worniu or En. (D) Schematic. Vnd and En are both required for Fd4 expression.

### Vnd misexpression generates ectopic Fd4+ ventral column NBs at the expense of Msh+ lateral NBs

Vnd is a known ventral column STF that promotes ventral column NB identity and represses lateral NB identity (McDonald et al., 1998), but in this older work the phenotypes were not assayed at the level of individual NB markers such as Fd4. To determine whether Vnd can induce the formation of ectopic Fd4+ NB7-1s, we used *en-gal4* to drive *UAS-vnd* across the En posterior row domain. In controls, there was a single, ventrally positioned Fd4+ NB7-1 (Figure 3A; quantified in 3C), whereas Vnd misexpression resulted in an average of four ectopic Fd4+ NB7-1s (Figure 3B, quantified in 3C). Note that the ectopic NB7-1s were restricted to the medio-lateral En domain). Next, we assayed for the ability of Vnd to repress lateral NB identity. In controls, the lateral NB marker Msh were expressed in a lateral band of NBs and progeny (Figure 3D), whereas Vnd misexpression resulted in a loss of the lateral Msh+ NBs and progeny (Figure 3E). Our results confirm prior findings that Vnd is sufficient to promote ectopic ventral NB identity (McDonald et al., 1998), and further show that this ectopic identity includes NB-specific identity markers.

**Figure 3.**
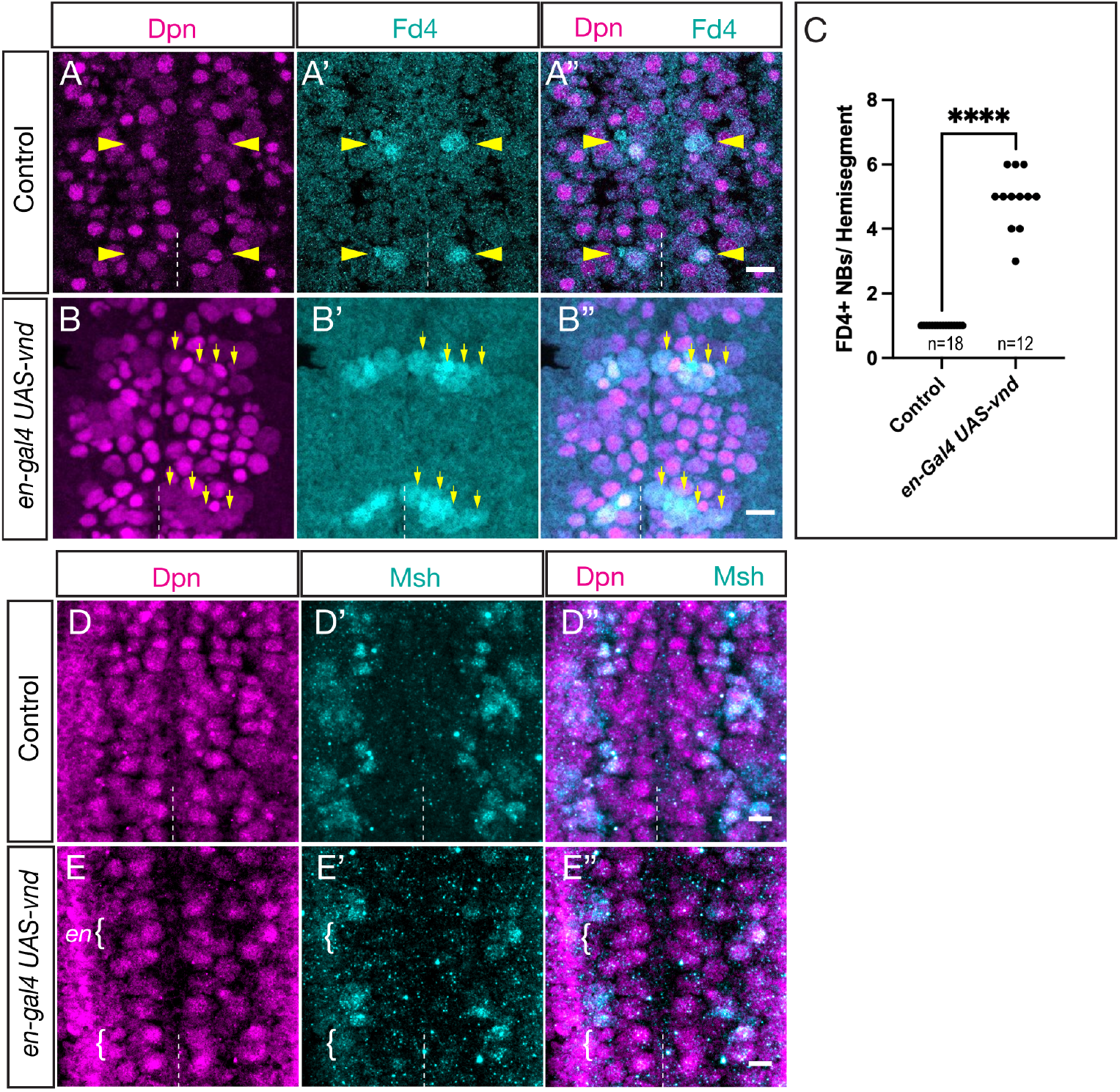
Vnd misexpression generates ectopic Fd4+ NBs at the expense of Msh+ lateral NBs. (A-C) Vnd misexpression increases Fd4+ NB numbers. (A-A’’) Control showing Fd4 expression in a single ventral column NB per hemisegment (yellow arrowhead). (B-B’’) Vnd misexpression in the En domain expands Fd4 into laterally located NBs. Dpn, pan-NB marker. Dashed line, ventral midline. Ventral view. Scale bar 10μm. (C) Quantification. (D-F) Vnd misexpression represses the lateral columnar marker Msh. (D-D’’) Control showing Msh expression in the lateral column of NBs. (E-E’’) Vnd misexpression in the En domain (brackets) represses Msh expression. Dashed line, ventral midline. Ventral view. Scale bar 10μm. (F) Quantification.

### Vnd misexpression generates ectopic NB7-1-derived neuronal progeny

We showed above that Vnd can promote Fd4+ NB7-1 identity; here we determined whether these ectopic NBs generated their normal Even-skipped (Eve)+ U1-U5 neuronal progeny. We used *en-gal4* to misexpress *UAS-vnd* and quantified the number of Eve+ U1-U5 neurons. In controls, there were invariantly five Eve+ U1-U5 neurons that could be identified using distinct molecular markers (see Methods) (Figure 4A; quantified in 4E, summarized in 4F). In contrast, Vnd misexpression generated ectopic U1-U5 neurons (Figure 4B; quantified in 4E, summarized in 4F); although it was difficult to identify individual U neurons, the total number is consistent with the average of ~4 more of each Fd4+ NBs described in the previous section. We conclude that Vnd misexpression can generate ectopic NB7-1s that undergo a normal cell lineage to generate ectopic U1-U5 neurons.

**Figure 4.**
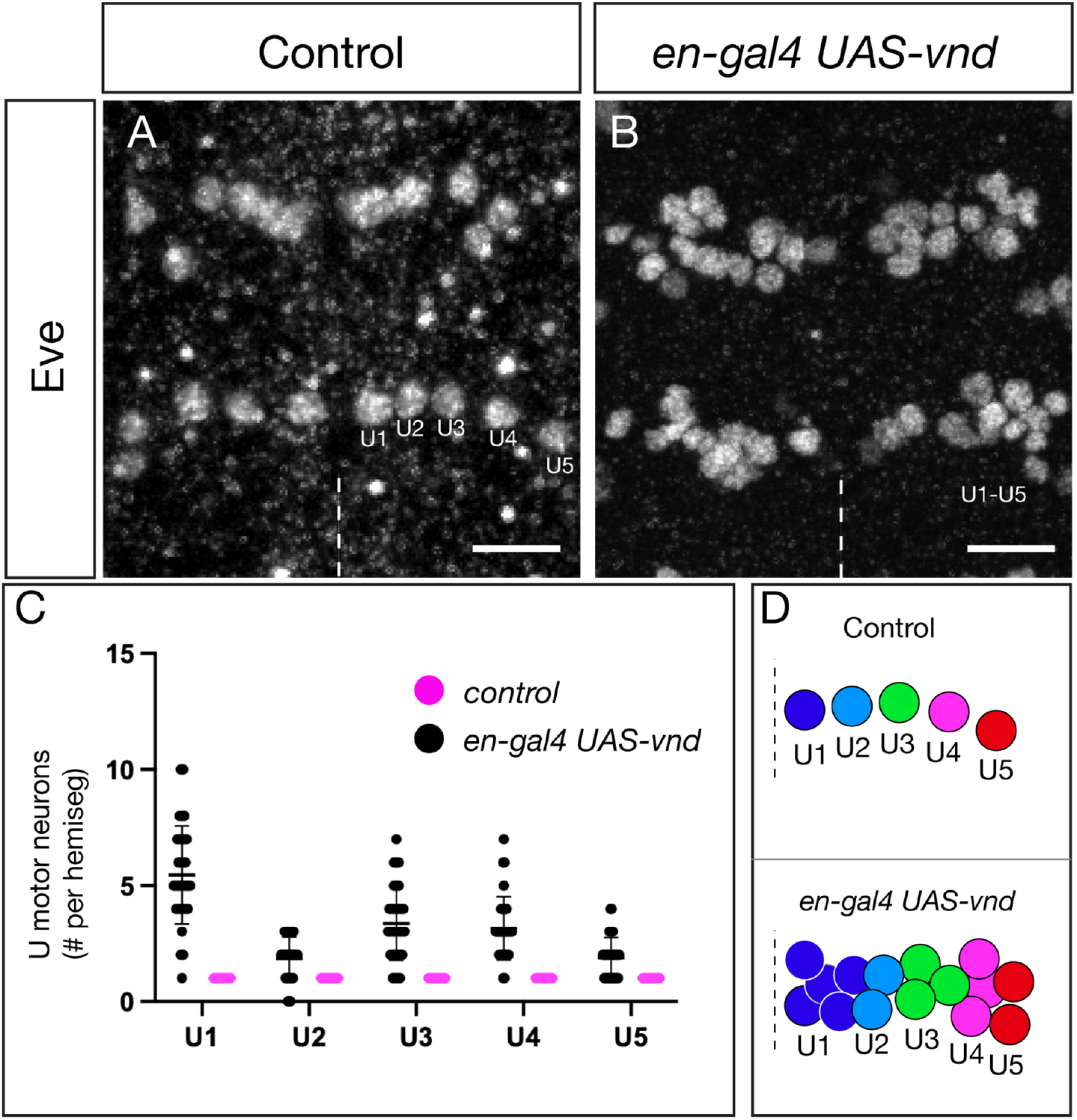
Vnd misexpression generates ectopic NB7-1-derived neuronal progeny. (A) Control showing five U1-U5 motor neurons from the NB7-1 lineage per hemisegment; two segments shown. U1 most medial, U5 most lateral positions. Dashed line, ventral midline; ventral view. Scale bar 10μm. (B) Vnd misexpression in the En domain results in ectopic NB7-1-derived U motor neurons. Two segments shown; dashed line, ventral midline; ventral view. Scale bar 10μm. (C) Quantification. (D) Schematic of phenotype.

### En misexpression generates ectopic Fd4+ NBs at the expense of row 5 Wg+ NBs

En is expressed in the posterior neuroectodermal rows (Chu-LaGraff and Doe, 1993; O’Farrell et al., 1985), making it an excellent candidate to be an STF. While it has been shown to be required for proper neurogenesis (Patel et al., 1989), it has never been tested for a role in specifying posterior row NB specification, and specifically in NB7-1 specification. To determine whether En can induce the formation of ectopic Fd4+ NBs, we generated a *vnd-gal4* transgenic line (see Methods) and used it to drive *UAS-en* across the anterior-posterior axis of the Vnd domain. In controls, there was a single, ventrally positioned Fd4+ NB (Figure 5A-A’’); in contrast, En misexpression in the Vnd expression domain resulted in two Fd4+ NBs, the endogenous NB7-1 and a more anterior ectopic Fd4+ NB (Figure 5B-B’’). The ectopic NB was restricted to the ventral, Vnd+ domain, consistent with both Vnd and En being required for specifying Fd4+ NB identity.

**Figure 5.**
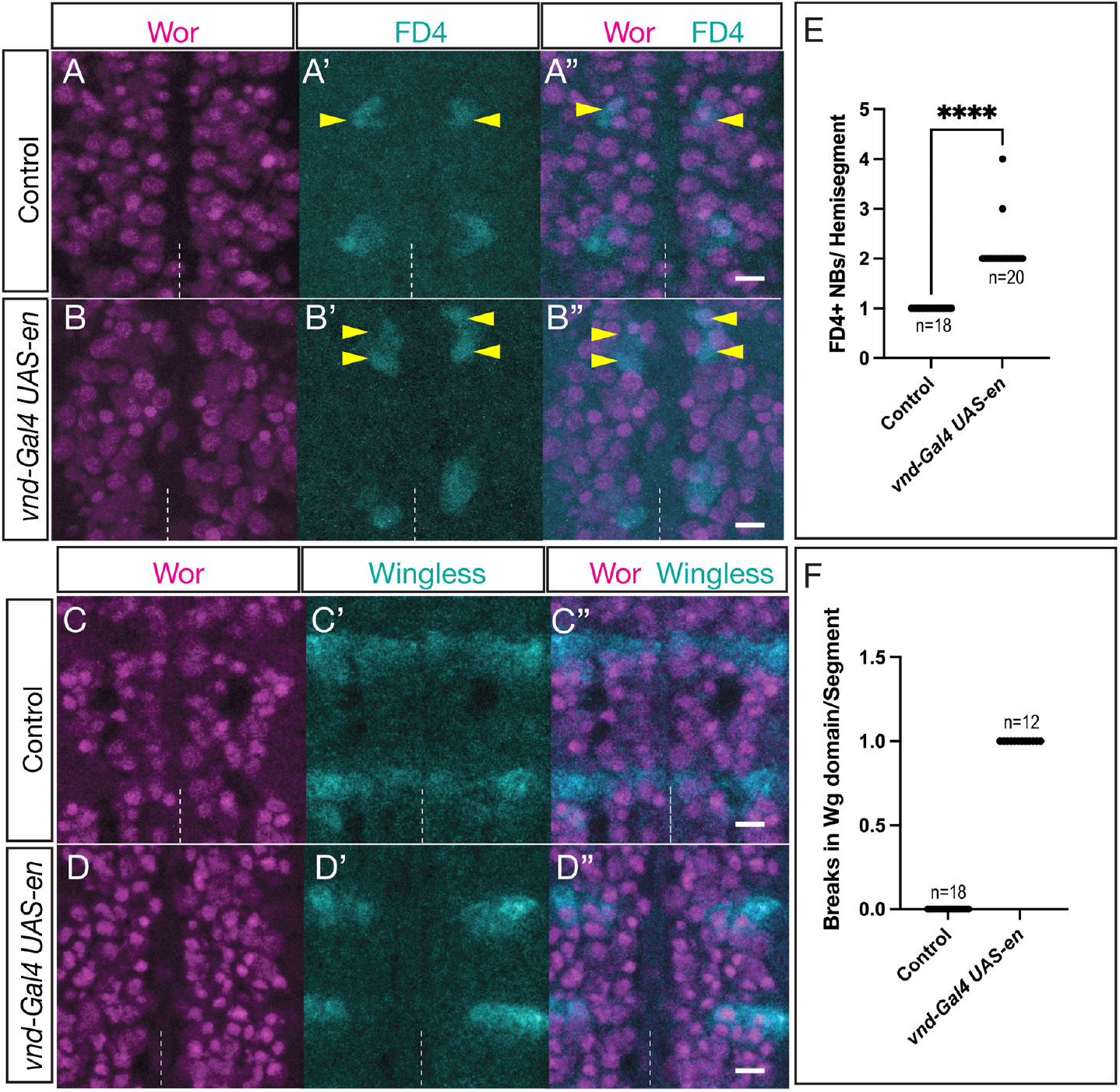
En misexpression generates ectopic Fd4+ NBs at the expense of Wg+ NBs. (A-A’’) Control (*vnd-gal4 UAS-lacZ*) showing one Fd4+ NB per hemisegment; Wor, pan-NB marker. (B-B’’) En misexpression in the ventral column of NBs (*vnd-gal4 UAS-en*) results in a single ectopic NB, resulting in two Fd4+ NBs per hemisegment (yellow arrowheads). Two segments shown. Stage 12; dashed line, ventral midline; ventral view. Scale bar 10μm. (C-C’’) Control (*vnd-gal4 UAS-lacZ*) showing expression of the row 5 marker Wingless. Stage 12; dashed line, ventral midline; ventral view. Scale bar 10μm. (D-D’’) En misexpression in the ventral column of NBs (*vnd-gal4 UAS-en*) results in repression of the row 5 marker Wingless. Stage 12; dashed line, ventral midline; ventral view. Scale bar 10μm.

Next, we asked: what NB identity was lost at the position of the ectopic NB7-1? We assayed Wingless, which was expressed in a row just anterior to NB7-1. In controls, Wingless was expressed in row 5 NBs across the entire medio-lateral axis of the segment, adjacent to the En+ rows 6/7 (Figure 5C-C’’). In contrast, En misexpression along the ventral column resulted in a loss of Wg expression in the ventral column, where NB5-1 and NB5-2 were located (Figure 5D-D’’). We conclude that En is sufficient to promote ectopic Fd4+ NB identity at the expense of the adjacent NB5-1 or NB5-2. It is not clear what prevents En from transforming the more anterior NBs in rows 1-4 (see Discussion).

### En misexpression generates ectopic NB7-1-derived neuronal progeny

We showed above that En misexpression could generate a single ectopic Fd4+ NB7-1. Here we determined whether the ectopic NB generates its normal, lineage-specific progeny. We used *vnd-gal4* to misexpress *UAS-en* and quantified the number of NB7-1-derived Eve+ U motor neurons. In controls, there were invariantly five Eve+ U1-U5 neurons (Figure 6A,C; quantified in 6E). In contrast, En misexpression generated ectopic Eve+ U neurons (Figure 6B,D; quantified in 6E). Note that due to the disruption in U neuron positions, we could only quantify the total number of Eve+ U neurons, and not individual U1-U5 neurons. Following En misexpression, there were ~2 fold the number of U neurons, consistent with the ~2 fold the number of Fd4+ NBs described in the previous section. We conclude that En misexpression can generate an ectopic NB7-1 that undergoes a normal cell lineage based on the production of five Eve+ U motor neurons.

**Figure 6.**
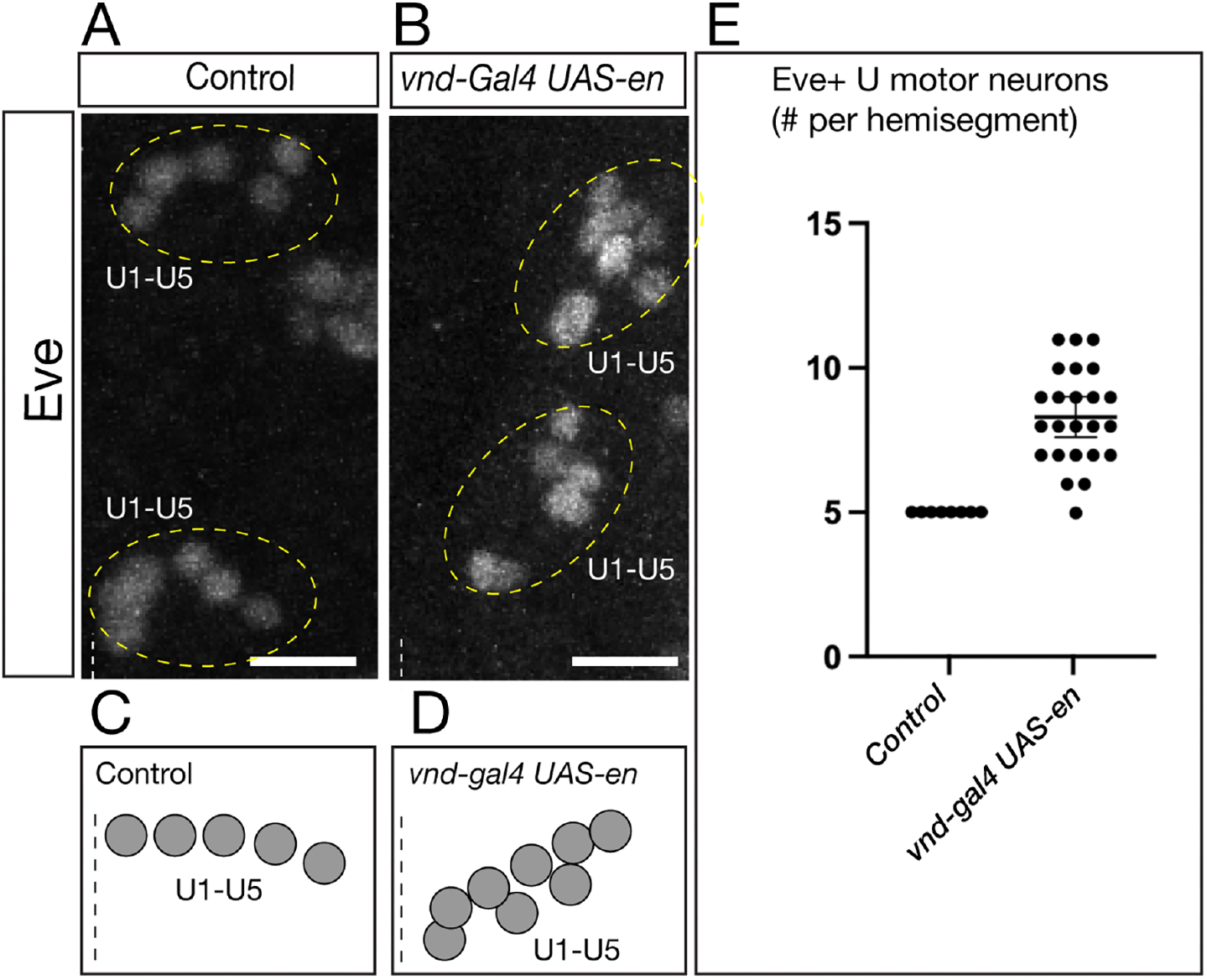
En misexpression generates ectopic NB7-1-derived neuronal progeny. (A) Control showing five Eve+ U neurons per hemisegment; two segments shown. Dashed line, ventral midline; ventral view. Anterior, up. Scale bar 10μm. (B) En misexpression in the Vnd domain results in ectopic Eve+ U neurons. Two segments shown; dashed line, ventral midline; ventral view. Anterior, up. Scale bar 10μm. (C,D) Schematic of phenotype. (E) Quantification. Scale bar 10μm.

### Neuroblasts gradually lose competence to respond to En and Vnd

We showed above that Vnd and En together specify NB7-1 identity, but it remains an open question when, or if, these NBs lose competence to respond to Vnd or En. Here we ask whether aging NBs lose competence to generate Fd4+ NB7-1 identity in respond to En and Vnd co-misexpression. To address this, we used three Gal4 lines expressed at progressively later times during neurogenesis: in the neuroectoderm (*sca-gal4*); in newly formed NBs (*wor-gal4*); and in NBs midway through their lineage (*grh-gal4*) (Figure 7A-E). In controls, we observed the expected single Fd4+ NB7-1 (Figure 7B-B’’). In contrast, *sca-gal4* induced Vnd and En co-misexpression in neuroectoderm showed increased numbers of Fd4+ NB7-1s (Figure 7C-C’’; quantified in F,G). Slightly fewer ectopic NB7-1s were generated by *wor-gal4*-induced Vnd and En co-misexpression in young NBs (Figure 7D-D’’; quantified in F,G), and an even smaller number of ectopic NB7-1s were induced by *grh-gal4* induced Vnd and En co-misexpression midway through NB lineages (Figure 7E-E’’; quantified in F,G). We conclude that NBs gradually lose competence to En and Vnd, but even mature NBs midway through their lineage retain some capacity for generating ectopic Fd4+ NB7-1s.

**Figure 7.**
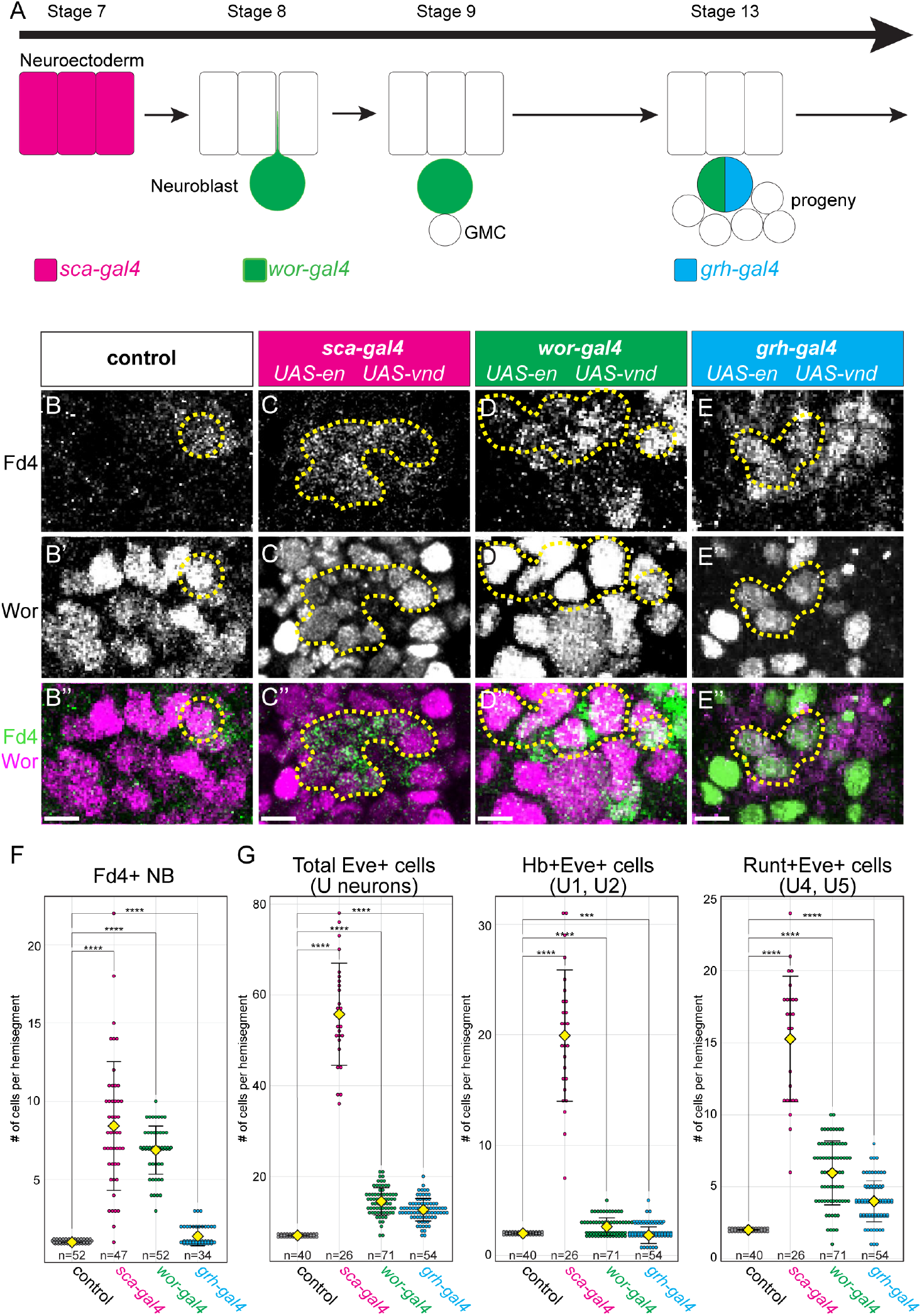
En and Vnd show progressive loss of competence to specify NB7-1. (A) Schematic showing distinct spatiotemporal expression of the indicated Gal4 transgenes. (B-B’’) Control; expression of indicated markers. (C-C’’) *sca-gal4 UAS-en UAS-vnd*; co-misexpression in the neuroectoderm results in ~7 FD4+ NBs. Dash lines circle the FD4+ NBs. Scale bar 10μm. (D-D’’) *wor-gal4 UAS-en UAS-vnd*; co-misexpression in the newly delaminated NBs results in ~7 FD4+ NBs. Dash lines circle the FD4+ NBs. Scale bar 10μm. (E-E’’) *grh-gal4 UAS-en UAS-vnd*; co-misexpression in the mid-lineage NBs results in ~3 FD4+ NBs. Dash lines circle the FD4+ NBs. Scale bar 10μm. (F-G) Quantification of FD4+ NBs (F) and Eve+ U neurons (G). Each dot represents a single hemisegmet. Yellow diamond, group average; error bars, standard deviation. Statistical analysis was performed using Student’s *t*-test. ***, p<0.001; ****, p<0.0001. Genotype: control, y-w-; sca-gal4, *sca-gal4 UAS-en UAS-vnd;* wor-gal4, *wor-gal4 UAS-en UAS-vnd*; grh-gal4, *grh-gal4 UAS-en UAS-vnd*.

## Discussion

In this work, we use the forkhead domain transcription factor Fd4 (Flybase: fd96Ca) as a marker for NB7-1 and its progeny. Within the CNS, Fd4 is specifically detected in NB7-1 and its lineage, although it has additional expression in leg imaginal discs (Heingård et al., 2019; Ruiz-Losada et al., 2021), embryonic peripheral nervous system (Häcker et al., 1992), and embryonic visceral mesoderm (Zhu et al., 2012). The detailed expression and functional analysis of Fd4 will be further analyzed in a separate paper (SLL and CQD, in preparation). Here we document its specific expression in the NB7-1 lineage and use it as a new marker for NB7-1.

Vnd is known to impart ventral column spatial identity in the neuroectoderm and NBs: loss of Vnd reduces ventral column markers and misexpression generates ectopic ventral column markers (McDonald et al., 1998). However, both loss-of-function and misexpression experiments focused on the role of Vnd in specifying neuroectoderm or NB identity, and only a few neurons were assayed (McDonald et al., 1998). Here we confirm these results and show that (1) Vnd misexpression generates ectopic Fd4+ NB7-1s at the expense of Msh+ lateral NBs; and (2) Vnd-induced ectopic NB7-1s undergo a normal cell lineage based on the production of U1-U5 motor neurons. The deviation from wild type is that the ectopic NB7-1s produce more early-born U neurons and less late-born U neurons. This may reflect the normal expression of Vnd in early-born neurons, followed by down-regulation at the time late-born neurons are produced. Moreover, we lack markers for the last-born neurons in the NB7-1 lineage (produced after U5) (Isshiki et al., 2001; Pearson and Doe, 2003), so we cannot say whether these last-born neurons are also made by ectopic NB7-1s.

En was previously shown to be required for normal embryonic CNS development (Bhat and Schedl, 1997; Patel et al., 1989). Neurons derived from the posterior of each segment are lost in the *en* mutants and there are duplications of neurons derived from anterior portions of each segment (Bhat and Schedl, 1997; Patel et al., 1989). However, NB identity was not monitored with NB-specific markers and En was not shown to act combinatorially to specify individual NB identities. Here we show that En is a spatial TF that acts together with Vnd to specify the ventral, posterior NB7-1. In addition, previous studies have only examined En loss-of-function, whereas we analyze both loss-of-function and misexpression experiments. Nevertheless, previous work and our work are entirely consistent and mutually supporting.

We show that En misexpression generates ectopic Fd4+ NB7-1s at the expense adjacent, row 5 Wg+ NBs. Interestingly, En misexpression in the ventral column (*vnd-Gal4 UAS-en*) results in only a single row 5 NB being transformed into an ectopic NB7-1. The anterior NB rows (1-4) may not have competence to respond to En misexpression, e.g. if En target genes are in silent chromatin domains. Alternately, there may be an En cofactor limited to rows 6/7 that is required for En to induce NB7-1 identity. The spatial TFs used to specify row 1-4 identity remain unknown.

We find that NBs gradually lose competence to respond to En and Vnd STFs; misexpression at progressively later times in neurogenesis shows a gradual decline in the number of ectopic NB7-1s that can be induced. This loss of responsiveness might be due to a general increase in genome-wide chromatin status (Sen et al., 2019) or more specific movement of En and Vnd target genes to the nuclear lamina or another repressive domain (Kohwi et al., 2013). Nevertheless, it is clear that some ectopic NB7-1s can be generated by late, mid-lineage expression to En or Vnd. It may be that En and Vnd act on the entire lineage to convert NB, GMC, and neurons into a NB7-1 lineage identity, or they may act on the NB only, giving rise to mixed lineages in which the first-born neurons are not converted but the later-born neurons are converted.

We note that the combination of Vnd and En is sufficient to specify NB7-1 identity; in the future it would be interesting to map En and Vnd binding sites at the *fd4* locus, which would help determine whether En and Vnd act directly to activate *fd4* transcription.

## Acknowledgments

We thank Tom Kornberg and Judith Kassis for *UAS-en* flies, Dervla Mellerick for *UAS-vnd* flies, Keiko Hirono for making *vnd-T2A-gal4* flies. We thank Kasey Drake, Derek Epiney and Keiko Hirono for comments on the manuscript. Antibodies obtained from the Developmental Studies Hybridoma Bank, created by the NICHD of the NIH and maintained at the University of Iowa, Department of Biology, Iowa City, IA were used in this study. Stocks obtained from the Bloomington Drosophila Stock Center (NIH P40OD018537). Funding was provided by NIH R01 HD27056 (CQD), HHMI (CQD and S-LL), and NIH Dev Biology Training Grant T32-ND007348 (NA).

This article is subject to HHMI’s Open Access to Publications policy. HHMI lab heads have previously granted a nonexclusive CC BY 4.0 license to the public and a sublicensable license to HHMI in their research articles. Pursuant to those licenses, the author-accepted manuscript of this article can be made freely available under a CC BY 4.0 license immediately upon publication.

## Author contributions

Conceptualization NA, SLL, CQD; Data curation NA, SLL, CQD; Formal analysis NA, SLL, CQD; Funding acquisition NA,CQD; Investigation NA, SLL; Methodology NA, SLL; Project administration CQD; Supervision CQD; Validation NA, SLL, CQD; Visualization NA, SLL, CQD; Roles/Writing - original draft NA, SLL, CQD; and Writing - review & editing NA, SLL, CQD.

## References

Albertson R, Doe CQ. 2003. Dlg, Scrib and Lgl regulate neuroblast cell size and mitotic spindle asymmetry. Nat Cell Biol 5:166–70.

Bhat KM. 1996. The patched signaling pathway mediates repression of gooseberry allowing neuroblast specification by wingless during Drosophila neurogenesis. Development 122:2921–32.

Bhat KM, Schedl P. 1997. Requirement for engrailed and invected genes reveals novel regulatory interactions between engrailed/invected, patched, gooseberry and wingless during Drosophila neurogenesis. Dev Camb Engl 124:1675–1688.

Briscoe J, Pierani A, Jessell TM, Ericson J. 2000. A homeodomain protein code specifies progenitor cell identity and neuronal fate in the ventral neural tube. Cell 101:435–445.

Cheesman SE, Layden MJ, Von Ohlen T, Doe CQ, Eisen JS. 2004. Zebrafish and fly Nkx6 proteins have similar CNS expression patterns and regulate motoneuron formation. Development 131:5221– 32.

Chu H, Parras C, White K, Jiménez F. 1998. Formation and specification of ventral neuroblasts is controlled by vnd in Drosophila neurogenesis. Genes Dev 12:3613–3624.

Chu-LaGraff Q, Doe CQ. 1993. Neuroblast specification and formation regulated by wingless in the Drosophila CNS. Science 261:1594–7.

Cleary MD, Doe CQ. 2006. Regulation of neuroblast competence: multiple temporal identity factors specify distinct neuronal fates within a single early competence window. Genes Dev 20:429–34.

Grosskortenhaus R, Pearson BJ, Marusich A, Doe CQ. 2005. Regulation of temporal identity transitions in Drosophila neuroblasts. Dev Cell 8:193–202. doi:10.1016/j.devcel.2004.11.019

Häcker U, Grossniklaus U, Gehring WJ, Jäckle H. 1992. Developmentally regulated Drosophila gene family encoding the fork head domain. Proc Natl Acad Sci U S A 89:8754–8758. doi:10.1073/pnas.89.18.8754

Heingård M, Turetzek N, Prpic N-M, Janssen R. 2019. FoxB, a new and highly conserved key factor in arthropod dorsal–ventral (DV) limb patterning. EvoDevo 10:28. doi:10.1186/s13227-019-0141-6

Isshiki T, Pearson B, Holbrook S, Doe CQ. 2001. Drosophila Neuroblasts Sequentially Express Transcription Factors which Specify the Temporal Identity of Their Neuronal Progeny. Cell 106:511–521. doi:10.1016/S0092-8674(01)00465-2

Kohwi M, Lupton JR, Lai SL, Miller MR, Doe CQ. 2013. Developmentally regulated subnuclear genome reorganization restricts neural progenitor competence in Drosophila. Cell 152:97–108. doi:10.1016/j.cell.2012.11.049

McDonald JA, Holbrook S, Isshiki T, Weiss J, Doe CQ, Mellerick DM. 1998. Dorsoventral patterning in the Drosophila central nervous system: the vnd homeobox gene specifies ventral column identity. Genes Dev 12:3603–12.

O’Farrell PH, Desplan C, DiNardo S, Kassis JA, Kuner JM, Sher E, Theis J, Wright D. 1985. Embryonic pattern in Drosophila: the spatial distribution and sequence-specific DNA binding of engrailed protein. Cold Spring Harb Symp Quant Biol 50:235–242. doi:10.1101/sqb.1985.050.01.030

Patel NH, Schafer B, Goodman CS, Holmgren R. 1989. The role of segment polarity genes during Drosophila neurogenesis. Genes Dev 3:890–904. doi:10.1101/gad.3.6.890

Pearson BJ, Doe CQ. 2003. Regulation of neuroblast competence in Drosophila. Nature 425:624–8.

Ruiz-Losada M, Pérez-Reyes C, Estella C. 2021. Role of the Forkhead Transcription Factors Fd4 and Fd5 During Drosophila Leg Development. Front Cell Dev Biol 9:723927. doi:10.3389/fcell.2021.723927

Sagner A, Zhang I, Watson T, Lazaro J, Melchionda M, Briscoe J. 2021. A shared transcriptional code orchestrates temporal patterning of the central nervous system. PLoS Biol 19:e3001450. doi:10.1371/journal.pbio.3001450

Schmid A, Chiba A, Doe CQ. 1999. Clonal analysis of Drosophila embryonic neuroblasts: neural cell types, axon projections and muscle targets. Development 126:4653–89.

Sen SQ, Chanchani S, Southall TD, Doe CQ. 2019. Neuroblast-specific open chromatin allows the temporal transcription factor, Hunchback, to bind neuroblast-specific loci. Elife 8. doi:10.7554/eLife.44036

Seroka A, Lai S-L, Doe CQ. 2022. Transcriptional profiling from whole embryos to single neuroblast lineages in Drosophila. Dev Biol 489:21–33.

Seroka AQ, Doe CQ. 2019. The Hunchback temporal transcription factor determines motor neuron axon and dendrite targeting in Drosophila. Development 137. doi:10.1242/dev.175570

Skeath JB, Carroll SB. 1992. Regulation of proneural gene expression and cell fate during neuroblast segregation in the Drosophila embryo. Development 114:939–46.

Skeath JB, Panganiban GF, Carroll SB. 1994. The ventral nervous system defective gene controls proneural gene expression at two distinct steps during neuroblast formation in Drosophila. Development 120:1517–1524. doi:10.1242/dev.120.6.1517

Skeath JB, Thor S. 2003. Genetic control of Drosophila nerve cord development. Curr Opin Neurobiol 13:8–15.

Skeath JB, Zhang Y, Holmgren R, Carroll SB, Doe CQ. 1995. Specification of neuroblast identity in the Drosophila embryonic central nervous system by gooseberry-distal. Nature 376:427–30.

Sullivan LF, Warren TL, Doe CQ. 2019. Temporal identity establishes columnar neuron morphology, connectivity, and function in a Drosophila navigation circuit. eLife 8. doi:10.7554/eLife.43482

Tabata T, Schwartz C, Gustavson E, Ali Z, Kornberg TB. 1995. Creating a Drosophila wing de novo, the role of engrailed, and the compartment border hypothesis. Dev Camb Engl 121:3359–3369. doi:10.1242/dev.121.10.3359

Tran KD, Doe CQ. 2008. Pdm and Castor close successive temporal identity windows in the NB3-1 lineage. Development 135:3491–9. doi:10.1242/dev.024349

Tran KD, Miller MR, Doe CQ. 2010. Recombineering Hunchback identifies two conserved domains required to maintain neuroblast competence and specify early-born neuronal identity. Development 137:1421–30. doi:10.1242/dev.048678

Weiss JB, Von Ohlen T, Mellerick DM, Dressler G, Doe CQ, Scott MP. 1998. Dorsoventral patterning in the Drosophila central nervous system: the intermediate neuroblasts defective homeobox gene specifies intermediate column identity. Genes Dev 12:3591–602. doi:PMID:%209832510

Zhu X, Ahmad SM, Aboukhalil A, Busser BW, Kim Y, Tansey TR, Haimovich A, Jeffries N, Bulyk ML, Michelson AM. 2012. Differential regulation of mesodermal gene expression by Drosophila cell type-specific Forkhead transcription factors. Development 139:1457–1466. doi:10.1242/dev.069005

